# Constitutive, endogenous, fluorescent membrane reporters for dynamic cell cycle analysis in *Bacillus subtilis*

**DOI:** 10.64898/2026.01.24.701471

**Authors:** Jane H. Joncha, Sadie Ruesewald, Kehinde O. Adebiyi, Daniel B. Kearns, Stephen C. Jacobson

## Abstract

Bacteria increase in biomass and divide, but determining precisely when cell division completes is technically challenging. To aid time-lapse imaging and cell-cycle tracking, we set out to identify a protein in *Bacillus subtilis*, which when fused with a fluorophore would cause the membrane to fluoresce in a manner that was constitutive, uniform, and bright. A forward genetic transposon-based approach combined with fluorescence-activated cell sorting was used to identify a fluorescent fusion to the glucose PTS transport transmembrane protein PtsG with all desired properties. Moreover, PtsG-GFP was constitutive and neutral to growth under all conditions tested and also labeled membranes during sporulation. We used PtsG-GFP to track cell growth in microfluidic channels and determine when cytokinesis occurred, defined as when fluorescence reached a local maximum at the division plane. Simultaneous imaging with a compatible fluorescent fusion to the cell division protein FtsZ indicated that FtsZ peak intensity occurred midway through septum constriction and that Z-ring recycling coincided with cytokinesis. We conclude that PtsG-GFP is a powerful tool for membrane imaging and cell cycle tracking. As such, we provide constructs with fluorophores that emit across the visible spectrum and antibiotic resistance cassettes to facilitate deployment in *B. subtilis*.

**IMPORTANCE:** Bacterial cells are fully divided when new membrane separates the cytoplasm of each daughter. Reproducibly staining of bacterial membranes with exogenous labels for fluorescence microscopy can be challenging, particularly during chemostatic growth in microfluidic devices. Here, we report that fusion of a fluorescent protein to the glucose transport protein PtsG causes the membrane of *Bacillus subtilis* to give off bright and even fluorescence under a variety of conditions. We use PtsG-GFP to operationally define when cytokinesis occurs during growth, and we note that a fluorescent PtsG fusion would likely make fluorescent staining of the membrane more facile theoretically in any organism.

## INTRODUCTION

Cellular growth occurs when cells coordinate an increase in biomass with division and is the most important, complex, and powerful phenotype of bacteria. Laboratory techniques for measuring bacterial growth each have their strengths and weaknesses. Spectrophotometry measures growth by light absorption of a culture and is facile but indirect, as it most directly measures biomass increase and not cell division per se. Viable counting involves dilution plating to track cell division events as a change in colony forming units, but the results lag the actual measurement as time is needed for cells to grow to form visible colonies. Both techniques are often performed on batch cultures, where growth rates change over time either due to the depletion of nutrients or the accumulation of metabolic waste. Microscopic analysis of bacterial growth in microfluidic channels allows direct observation of cell division in a controlled chemostatic environment but requires specialized equipment (1–3). Moreover, determining when an individual cell has completed division is remarkably challenging.

One type of cellular growth is binary fission, where a mother cell divides in the middle to produce two identical daughter cells. The process begins once the mother reaches a particular mass, and treadmilling protofilaments of the cell division protein FtsZ condense into a ring at or near the geometric midpoint of the cell (4–8). FtsZ continues to be recruited to the midcell until a threshold is reached at which point a membrane-associated complex of peptidoglycan synthesis machinery, called the divisome, is recruited and activated (9–11). Next, the membrane constricts, and the FtsZ ring contracts (6,12,13). Once the constricting membrane fuses into a septum, cytokinesis, or the separation of daughter cell cytoplasms, is complete. In many Gram-positive bacteria like *Bacillus subtilis*, daughter cell separation begins after the completion of the division septum such that one often cannot tell if or when cell division has occurred by phase contrast microscopy.

In prior microfluidic experiments with *B. subtilis*, our group used a conservative definition of cytokinesis based on a 20% reduction in the fluorescence intensity of a cytoplasmic protein (14). We recognized that defining cytokinesis by cytoplasmic fluorescence loss however, might be complicated by the small scale of bacteria, the resolution limit of fluorescence microscopy, and the fact that cell separation does not necessarily have to occur after cell division. Thus, we considered that cytokinesis could be more directly and accurately measured as an increase in membrane fluorescence during septum formation. As commonly used membrane stains such as FM4-64 and TMA-DPH are difficult to deliver evenly over long periods of time, we sought to generate a constitutively expressed membrane-anchored fluorescent protein. Preliminary reverse genetic approaches to make either natively expressed or inducible alleles of fluorescent fusions to the transmembrane segment of either SwrB (15) or EzrA (16) failed to label the membrane, for unknown reasons (data not shown).

Here, we take a forward genetic approach to find that fusion of a fluorescent protein to the PTS transport component PtsG is brightly fluorescent, constitutive, and evenly labels the cell membrane. Moreover, the PtsG fusion is not deleterious to growth in either a variety of batch media conditions or chemostatic microfluidic channels. Microfluidic monitoring of PtsG-GFP fluorescence permitted a modified definition of cytokinesis when the local fluorescence intensity of the invaginating membrane reached a maximum. When compared to the previous definition, we found that a 20% loss of cytoplasmic fluorescence precedes cytokinesis by a quarter of the cell cycle and, while effective, should be revised to a 40% decrease to coincide with septum formation. Membrane staining also refined the analysis of FtsZ dynamics indicating FtsZ-mNeongreen continues to intensify after membrane invagination is initiated to a maximum, after which FtsZ disassembles to extinction at or near the point of membrane transection. We conclude that PtsG-GFP is a powerful and versatile tool for membrane staining that improves the resolution of microscopic growth analyses.

## RESULTS

### A forward genetic approach to isolate GFP fusions with membrane localization

To identify a candidate fusion that provides intense, constitutive, and evenly distributed membrane fluorescence, cells were mutagenized with a transposon capable of generating 3’ translational fusions to green fluorescent protein (GFP) (**Fig 1A**). Briefly, a plasmid carrying the spectinomycin-selectable Tn*FLXgfp* transposon was transformed into wild type DK1042 cells and grown at room temperature for extra-chromosomal maintenance of the delivery vehicle.

**Figure 1:**
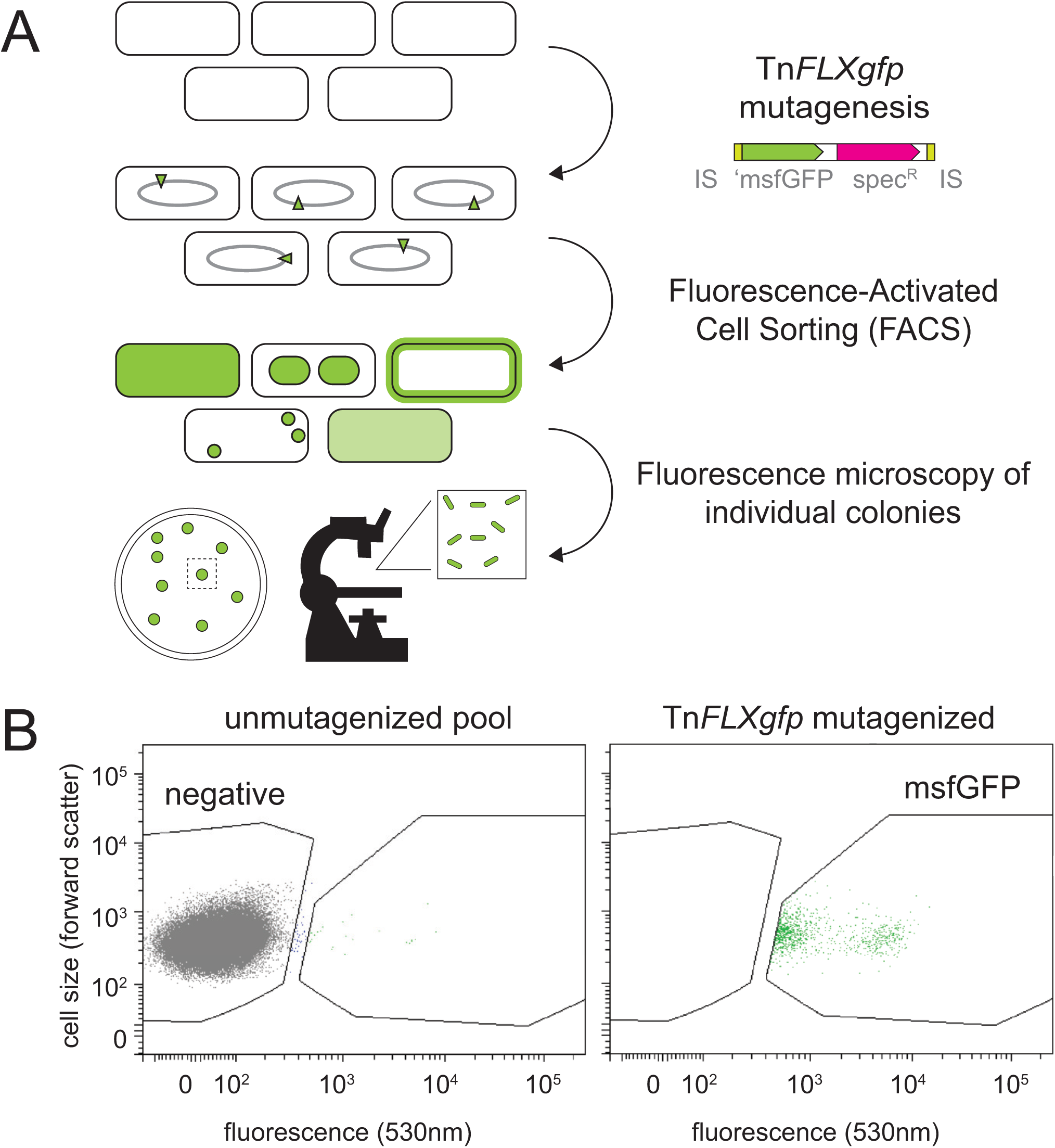
FACS-based strategy for isolating random translational fusions to GFP. A) A cartoon outline of the overall screening strategy. Wild type cells were mutagenized with a transposon carrying the spectinomycin antibiotic resistance cassette and an inward facing GFP gene lacking a ribosome binding site and translational start. IS elements indicated in yellow, msfGFP gene indicated as a green arrow, and spectinomycin resistance gene indicated as a magenta arrow. Thus, cells could only become fluorescent if the gene encoding GFP was inserted in frame into a gene that was being actively transcribed. To isolate cells containing fluorescent fusions, the pool was analyzed by a fluorescence activated cell sorter (FACS), and cells were selectively isolated if their fluorescence exceeded a threshold gate (**Fig S1**).Individual colonies were isolated from the pool of cells isolated for green fluorescence, manually grown, and observed by fluorescence microscopy. Finally, the location of each transposon was determined by inverse PCR and sequencing. B) FACS sorting data of the transposon mutagenized pool. Left indicates a FACS sort of unmutagenized wild type to establish the boundaries of non-fluorescent cells. Each gray dot represents a non-fluorescent individual. Right indicates a FACS sort of Tn*FLXgfp* mutagenized wild type. Each green dot indicates an individual with fluorescence in excess of a threshold gate. Note that there were also many non-fluorescent individuals in this population (gray circles) that were omitted due to an excess of datapoints. Ultimately, ∼2,000 green individuals were sorted from ∼40,000,000 cells in the total population so that candidates occurred at a frequency of 0.005%.

Colonies were picked into LB broth and grown to high density at 37°C, a temperature that is non-permissible for plasmid maintenance, and plated for high-density single colonies on plates containing spectinomycin, at the same temperature. The colonies were then pooled, back-diluted, grown to mid-exponential phase, and fluorescent individuals were separated by fluorescence activated cell sorting (FACS) (**Fig S1**). Specifically, all cells with fluorescence intensity more than a threshold established by comparison to a non-mutagenized, non-fluorescent control were pooled (**Fig 1B**) and dilution plated for single colony forming units. One hundred colonies were individually screened by fluorescence microscopy, and seventy were discarded as they failed to give detectable signal. The location of the transposon insertion was determined in the 30 remaining isolates, and after eliminating siblings, twelve strains were reserved for further analysis (**Table 1**).

**Table 1:**
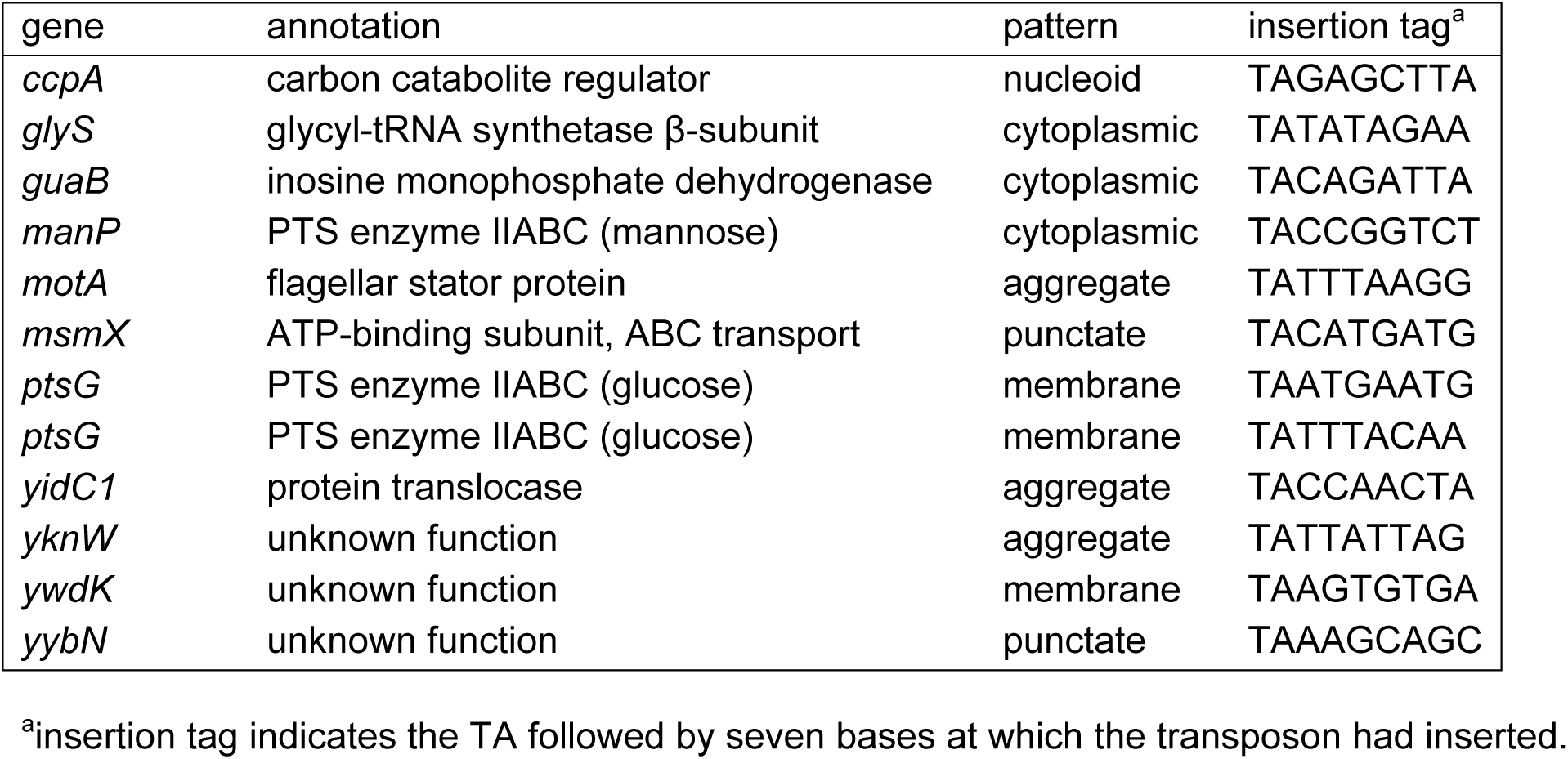
Genes in which *TnFLXgfp* had inserted and produced fluorescence.

Each of the unique insertions produced a variety of localization patterns in their respective strains. Three of the strains exhibited bright fluorescence that was diffuse in the cytoplasm (**Fig 2, cytoplasmic**). Two generated fusions to the predicted soluble proteins GlyS (glycyl-tRNA synthetase) and GuaB (inosine monophosphate dehydrogenase). The third was fused to the multi-pass transmembrane protein ManP (mannose phosphotransferase system) but was fused early in the protein prior to the first transmembrane segment. Three fusions appeared to form large aggregates and were fused to MotA (flagellar motor protein A), YidC1 (Sec-independent protein translocase), and YknW (unknown function). We infer that the aggregated localization pattern likely indicates dysfunctional fusions, as functional fusions to MotA appear as disparate puncta on the membrane (17), and the GFP fusion after a number of transmembrane segments in YidC1 might have been expected to have membrane localization (18–19).

**Figure 2:**
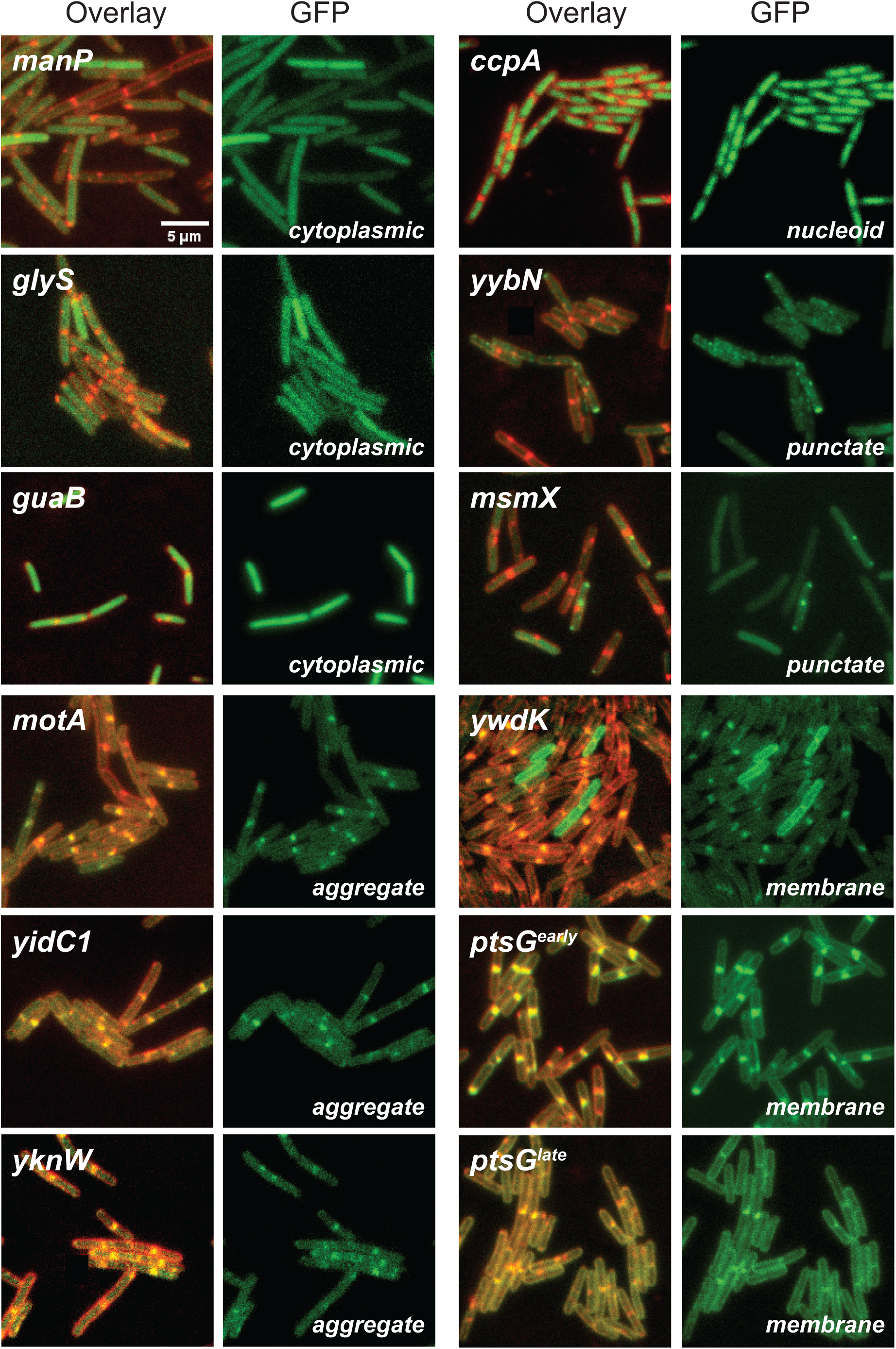
Cells expressing GFP fusions displayed a variety of subcellular localization patterns. Fluorescence micrographs of cells expressing a GFP fusion to the gene indicated in the upper left of each panel series. Left panel is an overlay of the membrane stain FM 4-64 (false colored red) and the fluorescence from the GFP fusion protein (false colored green). Each localization pattern was roughly characterized as cytoplasmic, aggregate, nucleoid, punctate, and membrane accordingly in the bottom right of each panel series (Table 1). The following strains were used for the generation of this figure: *manP* (DB1847), *glyS* (DB2138), *guaB* (DB2142), *motA* (DB2141), *yidC1* (DB2372), *yknW* (DB2136), *ccpA* (DB1701), *yybN* (DB1948), *msmX* (DB2140), *ywdK* (DB2137), *ptsG^early^* (DB1567), and *ptsG^late^* (DB1401). Scale bar is 5 μm.

One fusion in CcpA, the DNA-binding carbon catabolite regulator protein (20), had a localization pattern reminiscent of the nucleoid (**Fig 2, nucleoid**), and the fluorescence co-localized with DAPI staining (**Fig S2**). Two other fusions in YybN (unknown function) and MsmX (an ATP-dependent transport protein) (21,22) exhibited punctate localization patterns (**Fig 2, punctate**). Finally, three fusions appeared to give membrane localization (**Fig 2, membrane**). Fusion of GFP to YwdK (a protein of unknown function) strongly labeled the membrane but did so only in a minority subpopulation. The remaining two membrane-localized fusions were inserted at two locations in PtsG (enzyme IICBA of the glucose phosphotransferase system), one early in the coding sequence and one late. We conclude that the random Tn*FLXgfp* mutagenesis approach for generating fluorescent protein fusions was successful, and that the PtsG-GFP fusions were good candidates for a constitutive, endogenous membrane stain. We selected PtsG^early^-GFP (hereafter referred to as PtsG-GFP) for subsequent study as its fluorescence was brighter and more even than PtsG^late^-GFP.

### PtsG-GFP does not confer a growth defect

PtsG is a highly expressed membrane protein and mediates the high-affinity uptake of glucose (23–24). As the GFP fusion was integrated in, and thus, likely, disrupted the function of the native copy of the *ptsG* gene, we tested whether the PtsG-GFP fusion exhibited any pleiotropic effects on growth. Cells expressing PtsG-GFP grew like wild type in lysogeny broth (LB, rich undefined media), casein hydrolysate broth (CH, rich defined media), and S7_50_ broth (minimal defined media) supplemented with either 1% glucose or 1% glycerol (**Fig 3A**). We conclude that the expression of PtsG-GFP appeared to have little to no deleterious effects on cell growth. While we cannot account for all growth parameters, the absence of phenotype for those we tested support the use of PtsG-GFP as a neutral indicator of membrane localization in fluorescence microscopy. Moreover, while the operon encoding PtsG is poorly expressed in minimal media, fluorescence appeared constitutive and uniform under all growth conditions tested here perhaps because PtsG, which was inactivated by transposition, is required for autorepression (24).

**Figure 3:**
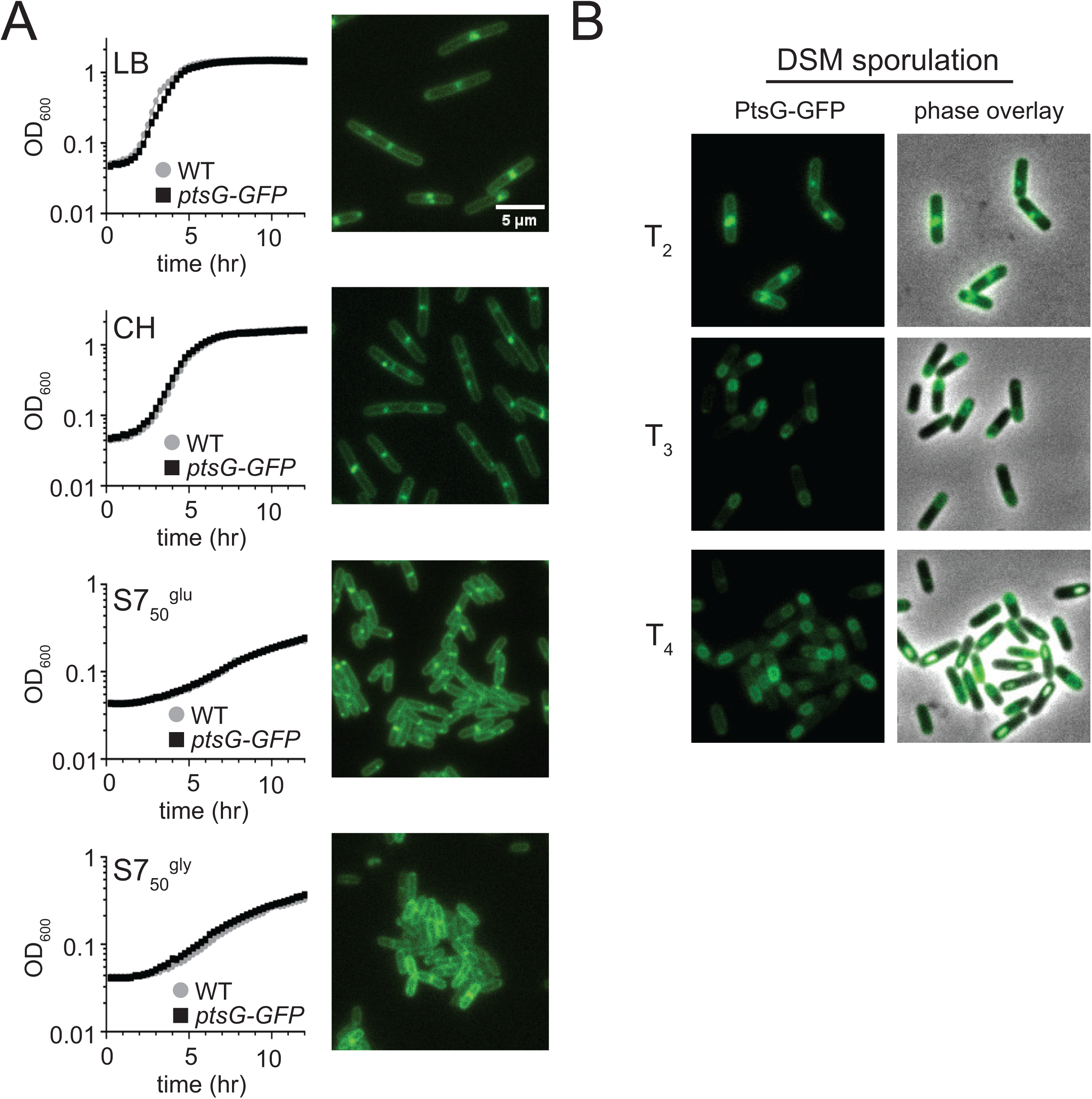
Cells expressing PtsG-GFP grow like wild type in a variety of media. (A) Graphs indicate growth curves conducted in plate reader with 1 ml volumes of the indicated media type and 24 well plates. Data are plotted as optical density at 600 nm (OD_600_) over time. Black squares, PtsG-GFP (DB1567); gray circles, wild type (DK1042). Micrographs indicate fluorescence of PtsG-GFP (DB1567) from cells grown in the same media as indicated by the graph at the left. Scale bar is 5 μm. (B) Micrographs false colored and overlayed shown under DSM sporulation indicate PtsG-GFP can track spore formation and indicate where spores have formed within the cell body. Micrographs were taken at hour 2, 3 and 4 after back dilution in DSM-complete media.

To deploy PtsG-GFP as a membrane reporter in a wide variety of genetic environments, we compared fluorescence in both the NCIB 3610 ancestral strain of *B. subtilis* as well as the commonly used laboratory strain derivative PY79. Fluorescence was strong in both ancestral and laboratory strain genetic backgrounds supporting general utility independent of genetic variation (**FIG S3A**). Moreover, fluorescence microscopy is often performed either by immobilizing cells on coverslips treated with poly-L-lysine, which while expedient can disrupt subcellular localization in some cases, or on agarose pads, which tends to preserve delicate subcellular localization patterns. Robust membrane staining was observed with both treatments, indicating that users could freely choose their mounting protocol depending on their protein of interest (**Fig S3B**). Finally, membrane fluorescence remained strong after the cessation of growth and induction of sporulation, suggesting that PtsG-GFP is also useful for labeling the membrane in post-exponential cells (**Fig 3B**).

To obtain higher resolution growth parameters at the level of individual cells, wild type and PtsG-GFP expressing cells were grown in microfluidic channels. For each strain, cell division events were measured with the previously established definition for cytokinesis of a 20% reduction in a constitutively expressed cytoplasmic mCherry signal (14). Wild type cells grew with a division time of 61 ± 12 minutes and an average cell length of 4.2 ± 0.9 μm. Cells expressing PtsG-GFP grew with a division time of 58 ± 18 minutes and an average cell length of 4.3 ± 0.9 μm. We note that the division time was longer than previously reported because the current measurements were taken at 30°C whereas previous publications grew cells at 37°C (14). Nonetheless, there was no statistically significant difference between division time or cell length between cells that expressed the PtsG-GFP fusion and cells that did not. We conclude that the presence of the PtsG-GFP fusion had no discernable effect on cell growth either in test tubes or in microfluidic devices.

### Endogenous membrane staining improves time resolution of cell division

The definition of cytokinesis based on a 20% reduction in cytoplasmic fluorescence was considered a conservative measurement of cell division because *B. subtilis* can complete septum formation before preceding to the conditional and non-essential step of cell separation (14,25,26). Given the problematic combination of the narrow septum (80 nm wide) (27), the resolution limit of fluorescence microscopy, and the fact that the measurement was a loss of signal, a 20% reduction was presumed to follow the event of septum formation by an indeterminate period. To determine whether PtsG-GFP could improve the time resolution of cytokinesis, the change in PtsG-GFP and cytoplasmic mCherry fluorescence intensity was compared over the course of the cell cycle in the same cells.

Cytoplasmic fluorescence was integrated along the cell body with a rectangular region of interest and expressed graphically as a line profile with periodic troughs (**Fig 4A**). The average cytoplasmic fluorescence intensity over multiple cell cycles exhibited a shallow negative slope (**Fig 4B, red**) with low dynamic range likely due to the nature of fluorescence loss and the point spread function of fluorescence affecting neighboring bins. PtsG-GFP expressed in the same cell, however, gradually increased in fluorescence intensity until it fully spanned the width and reached 100% signal intensity (**Fig 4A, 4B cyan star**). We conclude that defining cytokinesis as maximal local fluorescence of PtsG-GFP was more facile and provided a broader dynamic range than the loss of cytoplasmic fluorescence. Moreover, we note that contrary to expectations, a local 20% reduction in mCherry cytoplasm fluorescence (**Fig 4A, 4B red star**) preceded cytokinesis as determined by direct observation of septum formation and did so by approximately one quarter of the total cell cycle.

**Figure 4.**
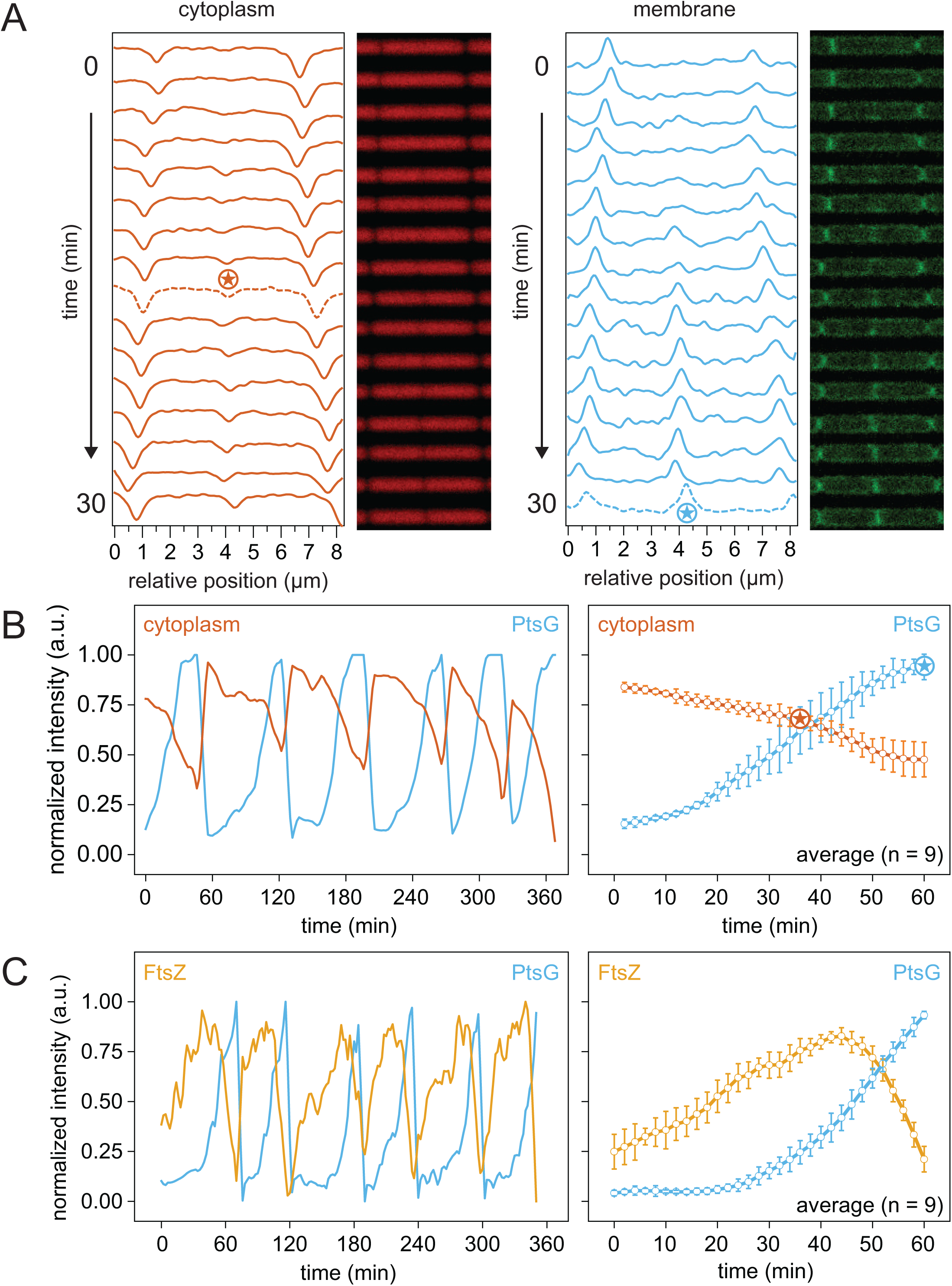
Cell biological comparison of cytoplasm labeling, membrane labeling, and mNeongreen-FtsZ. A) Fluorescence kymograph of strain DB1410 expressing cytoplasmic mCherry (false colored red, left) and PtsG-GFP (false colored green, right) during growth in a microfluidic channel. Each lane represents time points of growth separated by 2-min intervals. Graphs are fluorescence intensity traces of the micrograph lane immediately to the right. Red star indicates peak in the dotted red line where there was a local 20% reduction in cytoplasmic fluorescence intensity according to the definition of cytokinesis set by Yu et al., 2020. Cyan star indicates peak in the dotted cyan star where local membrane fluorescence intensity reached a maximum to create a new definition for cytokinesis. B) Left, fluorescence intensity of strain DB1410 expressing cytoplasmic mCherry (red), and the PtsG-GFP membrane label (cyan) of multiple cell division events expressed as arbitrary units (a.u.). Right, an average of fluorescence intensity of the cytoplasmic mCherry (red), and the PtsG-GFP membrane label (cyan) of n = 9 division events during growth in a microfluidic channel. Red star indicates point at which cytoplasmic fluorescence intensity decreased 20% and cyan star indicates point of maximum membrane intensity. Error bars are standard deviation. C) Left, fluorescence intensity of strain DB1526 expressing mCherry-FtsZ (gold), and the PtsG-GFP membrane label (cyan) of multiple cell division events expressed as arbitrary units (a.u.) during growth in a microfluidic channel. Right, average fluorescence intensity of mCherry-FtsZ (gold), and the PtsG-GFP membrane label (cyan) of n = 9 division events. Error bars are the standard deviation.

Previous work had indicated that the cell division protein FtsZ, persisted for a period of time at the cell pole after cytokinesis (14). The parameters of FtsZ dynamics, however, were measured in the context of using a reduction in cytoplasmic fluorescence that now seemed to substantially precede cytokinesis as measured by membrane staining. To study the relationship between FtsZ-ring dynamics and membrane synthesis, a strain expressing an FtsZ-mCherry fusion protein and a PtsG-GFP fusion simultaneously was grown in microfluidic devices. FtsZ fluorescence appeared first at the future site of cell division and increased over time (**Fig 4C**).

Indeed, FtsZ continued to accumulate at the midcell prior to the first appearance of membrane invagination presumably due to the maturation of the FtsZ ring that is needed to initiate septum construction (**Fig 4C**). After septum initiation, fluorescence intensity of both signals increased until the membrane fluorescence reached approximately 50% of maximum intensity, at which point FtsZ intensity peaked and began to decrease (**Fig 4C**). We conclude that FtsZ fluorescence intensity poorly correlates with that of the membrane, that peak FtsZ intensity occurs after the initiation of septum formation, and that FtsZ-ring disassembly is complete at or near the point of cytokinesis.

## DISCUSSION

The use of fluorescence microscopy for the cell biological analysis of protein function is common in *B. subtilis*, and membrane stains such as FM 4-64 or TMA-DPH are often used to provide context for protein subcellular localization. Fluorescent dyes are relatively inexpensive and easy to add for single-time-point analysis, but the staining procedure can be difficult to perfect, and excess dye incorporation may cause membrane perturbations. The challenge of consistent staining is exacerbated in experiments involving time-dependent observation. We sought to replace dye addition by the expression of a fluorescent protein that localizes to the membrane and is continually produced by *B. subtilis*. Here, we use an unbiased forward genetic approach to create random fluorescent protein fusions and screen for those that enrich in the membrane. We show that fluorescent protein fusions to PtsG provide for constitutive and intense membrane fluorescence suitable for both instantaneous and time-dependent observation over a variety of conditions without conferring a growth defect.

PtsG is the transmembrane component of the high-affinity phosphoenolpyruvate-dependent PTS transporter for glucose, and the native-site fusion used here is likely non-functional for glucose import. We were initially concerned that the fusion might cause growth defects but cells containing the PtsG-GFP fusion grew like wild type even when grown in minimal media with glucose as the sole carbon and energy source, presumably due to the presence of other glucose uptake systems. We were also concerned that PtsG-GFP fluorescence might be conditional as *ptsG* gene expression has been shown to vary (24), but the fluorescence intensity of PtsG-GFP seemed constant during both growth and sporulation. Satisfied that PtsG-GFP was both neutral and constitutive, four different fluorescent proteins were recombined with four different antibiotic cassettes to enhance compatibility with pre-existing cell biological constructs (**Table 2**, **Fig 5**). While native site integration restricts use to *B. subtilis* strains with high sequence similarity surrounding the *ptsG* locus, we note that PtsG is highly conserved across bacteria and suggest it would be a suitable candidate for generating constitutive membrane fluorescence in other organisms.

**Figure 5.**
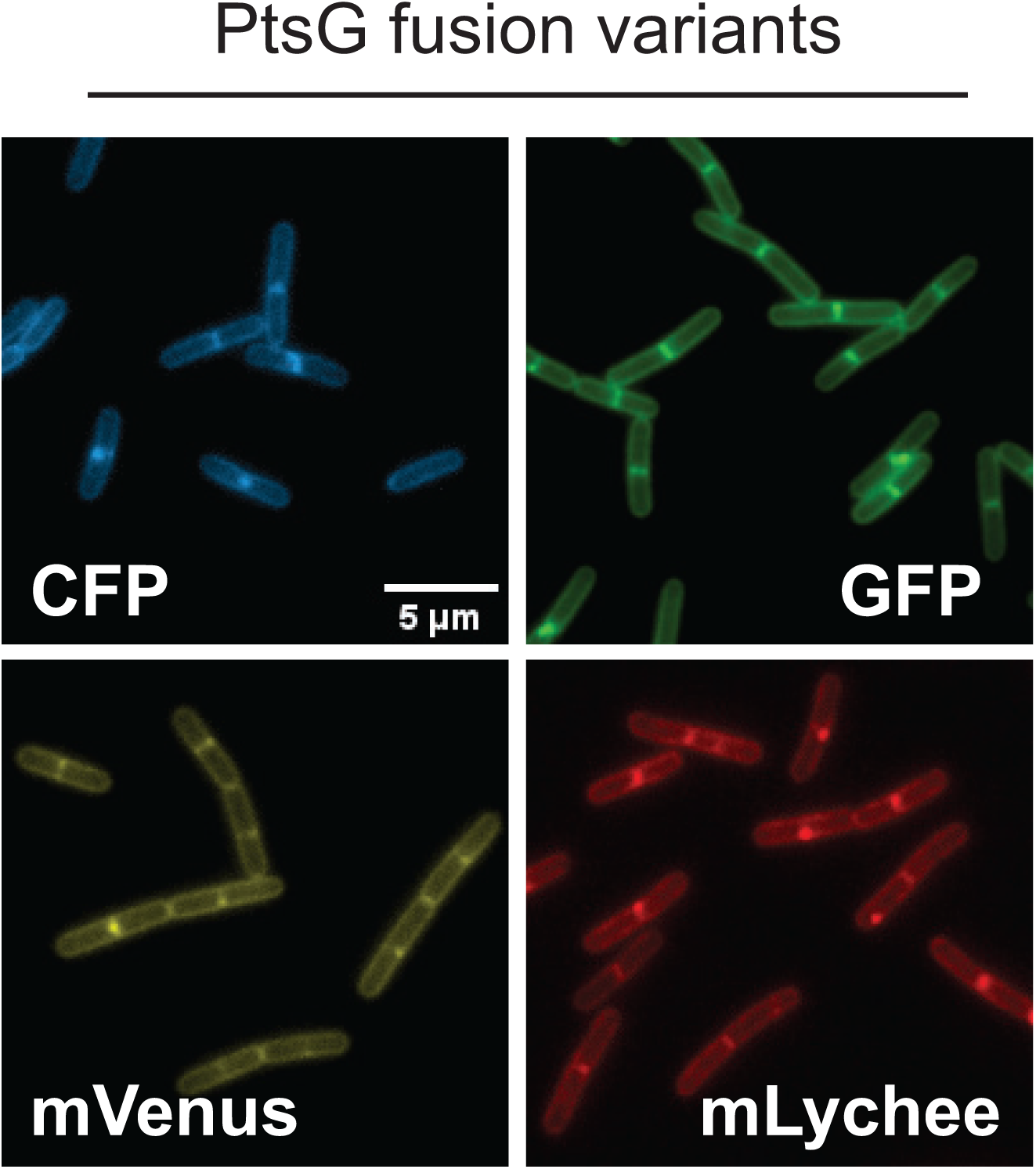
Variants of PtsG membrane labels with different fluorophore and antibiotic combinations. Fluorescent micrographs of the following strains: CFP (DB3548), GFP (DB3267), mVenus (DB3547), and mLychee (DB3500). Cells were imaged on 1% agarose pads, and the scale bar is 5 µm.

**Table 2:**
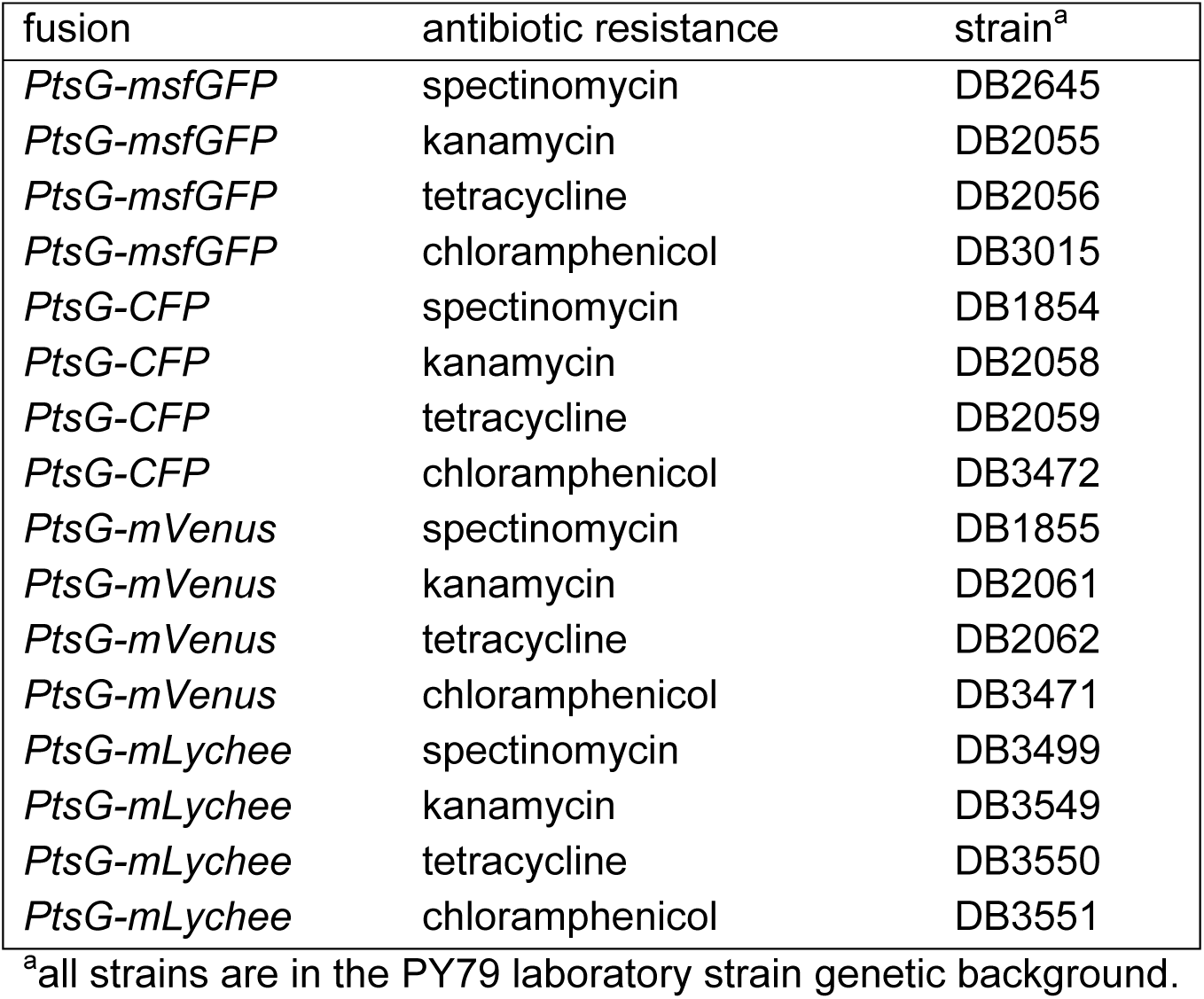
PtsG-fluorescent protein fusion constructs.

One application of PtsG-GFP is dynamic membrane labeling during microfluidic growth, and we suggest that the point at which septal membrane fluorescence reaches a maximum serves as the most direct and accurate definition of cytokinesis. By comparison, we find that cytokinesis defined by a 20% reduction in fluorescence of a cytoplasmically expressed protein precedes membrane transection by approximately one quarter of the cell cycle. Tracking the cytoplasm was nonetheless robust and changing to a 40% reduction in fluorescence would make it equivalent to the membrane-based definition. Simultaneous observation with a fluorescent FtsZ fusion indicates that FtsZ appears at the new division site before an increase in membrane fluorescence. FtsZ foci continue to intensify during membrane constriction to a maximum after which disassembly is rapid to extinction that roughly coincides with the formation of a membrane transect. Thus, we infer that the rate of FtsZ recruitment may be constant, but that the rate of disassembly either gradually increases during constriction or initiates after a checkpoint. Whatever the case, direct observation of membrane constriction clarified the order of events during division, and contrary to our conclusions using the cytoplasmic definition of cytokinesis (14), FtsZ does not appreciably dwell at the cell poles after division is complete, at least in the wild type. We conclude that PtsG-GFP is a robust tool for fluorescent labeling of the membrane and can be modified as necessary to meet the needs of the user.

Fluorescent fusions to PtsG recombining four different fluorophores and four different antibiotic resistance cassettes will be donated to the Bacillus Genetic Stock Center (BGSC, The Ohio State University) and the sequence of the PtsG-GFP is included in the supplemental materials to aid future modification.

## METHODS

### Strains and growth conditions

*B. subtilis* strains were grown in LB lysogeny broth (10 g tryptone, 5 g yeast extract, 5 g NaCl per L) or on LB plates fortified with 1.5% Bacto agar at 37°C. When appropriate, antibiotics were included at the following concentrations: 10 µg/ml tetracycline, 5 µg/ml chloramphenicol, 5 µg/ml kanamycin, 100 µg/ml spectinomycin, and 1 µg/ml erythromycin plus 25 µg/ml lincomycin (mls). Isopropyl β-D-thiogalactopyranoside (IPTG, Sigma) was added to the medium at the indicated concentration when appropriate. As needed, cells were grown in the rich defined media casein hydrolysate (CH), prepared as described previously (Harwood, 1990).

S7_50_ media was prepared in 100 mL aliquots, by mixing 10 mL 10X S7_50_ salts (recipe below), 1 mL 100X S7_50_ metals (recipe below), 2 mL 1 M glutamate and 2 mL of 50% glucose or glycerol together with 82 mL dH_2_O. All components (save for the dH_2_O) were filter sterilized.

10X S7_50_ salts were made in 1 L aliquots: 104.7 g MOPS (free acid), 13.2 g ammonium sulfate (NH_4_)2SO_4_, 6.8 grams potassium phosphate monobasic KH_2_PO_4_ were added and buffered to pH 7 with 1 M KOH in a volume of 400 mL of ddH_2_O, then made up to 1 L with ddH_2_O. Media was filter-sterilized into 100 mL aliquots, covered with foil, and stored at 4 °C. If yellowed, the solution is no longer usable. 100X S7_50_ metals have final concentrations of 0.2 M MgCl_2_, 70 mM CaCl_2_, 5 mM MnCl_2_, 0.1 mM ZnCl_2_, 2 µg/mL thiamine-HCl, 2 mM HCL, and 0.5 mM FeCl_3_. Fe was added last to prevent precipitation. Solution was filter sterilized and stored foil-wrapped in 10 mL aliquots at 4 °C.

DSM-incomplete broth was prepared as a 1 L bulk and subsequently dispensed as 100 mL aliquots. The 1 L DSM-incomplete broth was prepared by mixing 8 g of Difco Nutrient Broth, 10 mL of 100 mM MgSO_4_, 10 mL od 10% KCl, 0.5 mL of 1N NaOH and 1 L of dH_2_O. The 100 mL aliquots are then autoclaved for 30 minutes and stored at bench top conditions until ready for use. Once ready for use, add 100 µL of 1 M Ca(NO_3_)_2_, 100 µL of MnCl_2_, and 100 µL of 1 mM FeSO_4_ into a 100 mL DSM-incomplete broth aliquot to make 100 mL of DSMcomplete broth.

### Growth curve

Cells were grown in 3 ml of the desired media rolling at 37°C until turbid. Cultures were then back diluted to a starting OD_600_ of 0.01. On a 24 well plate, 500 µl was transferred into the wells in triplicate and then placed without the lid into the plate reader (Agilent BioTek synergy H1 microplate reader). The plate reader was set to a double orbital shaking on high at 37°C for 18 hr taking OD_600_ measurements every 15 min with Gen5 Microplate Reader and Imager Software Version 3.11. Data were collected the next day and graphed with GraphPad Prism 10 software.

### Preparation of cells for microscopy

Cells were grown in LB broth at 37°C until log phase (OD_600_ 0.5–0.9). A total of 1 mL of cells was then pelleted. Cells were resuspended in 30 µL 5 µg/mL FM4-64 (Molecular Probes) and incubated at room temperature for 3 min to stain the membrane before being pelleted again. Cells were then washed with 1 mL 1X PBS and pelleted again. All samples were resuspended in 30 µL 1X PBS, then observed by spotting 4 µL of this suspension on a 1% agarose pad made with water. CH media can sustain growth while maintaining low background fluorescence during imaging.

For sporulation, cells were grown in 2 mL of DSMcomplete broth for 1-2 hours at 37°C, then back diluted with 25 mL of DSMcomplete to an OD_600_ of 0.01 at 37°C. 1 mL of culture was collected every hour after the end of exponential phase to observe cell sporulation. All samples were centrifuged at 8000 x g for 1 min and supernatant was removed. Samples were then resuspended in 30 µL of dH_2_O and imaged.

### Microscopy

Fluorescence microscopy was performed with a Nikon 80i microscope along with a phase contrast objective Nikon Plan Apo 100x and an Excite 120 metal halide lamp. FM4-64 fluorescence signals were visualized with a C-FL HYQ Texas Red filter cube (excitation filter 532 to 587 nm, barrier filter 590 nm). Green fluorescent protein (GFP) was visualized with a C-FL HYQ FITC filter cube (fluorescein isothiocyanate [FITC], excitation filter 460 to 500 nm, bandpass filter 515 to 550 nm). Yellow fluorescent protein (YFP) was visualized with a C_FL HYQ YFP filter cube (excitation filter 490 to 510 nm, barrier filter 515 to 550 nm). TMA-DPH fluorescent signal was visualized with a UB-2E/C DAPI filter cube (excitation filter 340 to 380 nm, bandpass filter 435 to 485 nm). Images were captured with a Photometrics Coolsnap HQ2 camera in black and white, false-colored, and superimposed with NIS elements image software.

Fluorescence microscopy was also performed with a Nikon Ti2 Eclipse along with a phase contrast objective Nikon Plan Apo 100X oil-immersion objective and Lumencor spectra light. GFP was excited with 474/27 nm and collected at 515/30 nm. CFP was excited with 438/24 nm and collected at 483/32 nm. mVenus was excited with 509/22 nm and collected at 543/22 nm. mLychee signals were excited with 554/23 nm and collected at 595/31 nm. Semrock Triple Band Pinkel multiband and Quad band Sedat multiband filter sets were coupled with single band emission filters for generic fluorophore spectra. Images were captured with a Prime 95B Photometrics camera engine.

### Strain construction

Strain constructs were generated either by direct transformation into PY79 and transduced into 3610 by the generalized transducing phage SPP1 (28) or by direct transformation into the competent 3610 background strain DK1042 (29). All strains used in this study are listed in **Table 3**. All plasmids used in this study are listed in **Supplemental Table S1**. All primers used in this study are listed in **Supplemental Table S2**.

**Table 3:** Strains.

| strain | genotype |
| --- | --- |
| DK1042 | *comI*<sup>Q12L</sup> (ancestral background) |
| PY79 | *sfp*<sup>0</sup> *swrA*<sup>FS</sup> (laboratory strain) |
| DB1396 | [PY79] *ptsG*<sup>late</sup> ΩTnFLXgfp spec |
| DB1401 | *comI*<sup>Q12L</sup> *ptsG*<sup>late</sup> ΩTnFLXgfp spec |
| DB1526 | Δ*epsH* *yuxG*::*P*<sub>ftsA</sub>-mcherry-ftsZ mls *ptsG*ΩTnFLXgfp spec |
| DB1567 | *comI*<sup>Q12L</sup> *ptsG*<sup>early</sup> ΩTnFLXgfp spec |
| DB1701 | *comI*<sup>Q12L</sup> *ccpA*ΩTnFLXgfp spec |
| DB1756 | Δ*epsH* *amyE*::*P*<sub>hyspank</sub>-mcherry kan *ptsG*ΩTnFLXgfp spec |
| DB1847 | *comI*<sup>Q12L</sup> *manP*ΩTnFLXgfp spec |
| DB1854 | [PY79] *ptsG*ΩTnFLXcfp spec |
| DB1855 | [PY79] *ptsG*ΩTnFLXmvenus spec |
| DB1864 | *comI*<sup>Q12L</sup> *ptsG*ΩTnFLXcfp spec |
| DB1865 | *comI*<sup>Q12L</sup> *ptsG*ΩTnFLXmvenus spec |
| DB1948 | *comI*<sup>Q12L</sup> *yybN*ΩTnFLXgfp spec |
| DB2055 | [PY79] *ptsG*ΩTnFLXgfp kan |
| DB2056 | [PY79] *ptsG*ΩTnFLXgfp tet |
| DB2058 | [PY79] *ptsG*ΩTnFLXcfp kan |
| DB2059 | [PY79] *ptsG*ΩTnFLXcfp tet |
| DB2061 | [PY79] *ptsG*ΩTnFLXmvenus kan |
| DB2062 | [PY79] *ptsG*ΩTnFLXmvenus tet |
| DB2136 | *comI*<sup>Q12L</sup> *yknW*ΩTnFLXgfp spec |
| DB2137 | *comI*<sup>Q12L</sup> *ywdK*ΩTnFLXgfp spec |
| DB2138 | *comI*<sup>Q12L</sup> *glyS*ΩTnFLXgfp spec |
| DB2140 | *comI*<sup>Q12L</sup> *msmX*ΩTnFLXgfp spec |
| DB2141 | *comI*<sup>Q12L</sup> *motA*ΩTnFLXgfp spec |
| DB2142 | *comI*<sup>Q12L</sup> *guaB*ΩTnFLXgfp spec |
| DB2372 | *comI*<sup>Q12L</sup> *yidC1*ΩTnFLXgfp spec |
| DB2645 | [PY79] *ptsG*ΩTnFLXgfp spec |
| DB3015 | [PY79] *ptsG*ΩTnFLXgfp cat |
| DB3471 | [PY79] *ptsG*ΩTnFLXmvenus cat |
| DB3472 | [PY79] *ptsG*ΩTnFLXcfp cat |
| DB3499 | [PY79] *ptsG*ΩTnFLXmlychee spec |
| DB3549 | [PY79] *ptsG*ΩTnFLXmlychee kan |
| DB3550 | [PY79] *ptsG*ΩTnFLXmlychee tet |
| DB3551 | [PY79] *ptsG*ΩTnFLXmlychee cat |
| DB3267 | *comI*<sup>Q12L</sup> *ptsG*-cfp tet |
| DB3500 | *comI*<sup>Q12L</sup> *ptsG*-gfp cat |
| DB3547 | *comI*<sup>Q12L</sup> *ptsG*-mvenus kan |
| DB3548 | *comI*<sup>Q12L</sup> *ptsG*-mlychee spec |
| DK5133 | Δ*epsH* mNeongreen-ftsZ *amyE*::*P*<sub>hyspank</sub>-mcherry spec |
| DK9790 | *yuxG*::*P*<sub>ftsA</sub>-ftsZ mls |

### FtsZ-mCherry

To generate pLL4, the promoter region of *ftsA* was amplified from DK1042 chromosomal DNA with primers 7611/7606 and digested with HindIII/NheI, and the *ftsZ* gene was amplified from DK1042 chromosomal DNA with primers 7607/7608 and digested with NheI/BamHI. The two fragments were cloned simultaneously into the HindIII/BamHI sites of pWX146 containing a polylinker and erythromycin resistance cassette between two arms of the *yuxG* gene (generous gift of Dr. Xindan Wang, Indiana University). Plasmid pWX146 was then transformed into DK1042 to generate strain DK9790.

To generate an mCherry fusion to FtsZ, the region upstream of *yuxG* including the promoter region of *ftsA* was amplified from DK9790 chromosomal DNA with primers 7758/7759, the ftsZ gene, erythromycin resistance cassette, and region downstream of *yuxG* was amplified from DK9790 chromosomal DNA with primers 7762/7763, and the mCherry gene was amplified from pDR201 with primers 7760/7761 (generous gift of Dr. David Rudner, Harvard Medical School). The three amplicons were stitched together with isothermal assembly (30) and transformed into DK1042.

### Fluorophore replacements

To generate the PtsG-spec alleles containing different fluorophores such as cyan fluorescent protein (CFP), mVenus and mLychee, each cassette was separately PCR amplified from either pDR200 (*cfp*), pFK116 (*mvenus*) or pDP679 (*mlychee*).

The primer pairs for each plasmid were 8498/8499 for *cfp*, 8488/8489 for *mvenus* and 9060/9061 for *mlychee* (31). Two fragments, one containing the 5’ end of the *ptsGΩpTnFLXgfp* insertion plus the region upstream, and the other containing the 3’ end of the *ptsGΩpTnFLXgfp* insertion plus the region downstream, were separately PCR amplified from DB1567 chromosomal DNA using primers specific to each fluorophore: *cfp* (8486/8490 and 8491/8487), *mvenus* (8486/8496 and 8491/8487), and *mlychee* (8486/8886 and 9062/8487). Each of the two chromosomally amplified fragments was mixed with a fragment encoding the cognate fluorophore cassette and ligated by Gibson isothermal assembly (30). The resulting fragments were transformed into PY79 selecting spectinomycin resistance. The constructs were confirmed by amplifying the region using primer pair 8486/8487 and sequencing the amplicon.

### Antibiotic replacements

To generate the PtsG-GFP alleles containing kanamycin (kan) and tetracycline (tet) resistance cassettes, each cassette was separately amplified from either pDG780 (kan) or pDG1515 (tet) (32) with primer pair 3250/3251. Two other fragments containing the 5’ *ptsGΩpTnFLXgfp* insertion plus the region upstream, and a fragment containing the 3’ end of the *ptsGΩpTnFLXgfp* insertion plus the region downstream, were separately PCR amplified from DB1567 chromosomal DNA with primer pairs 8486/8516 and 8487/8517. Each of the two chromosomally amplified fragments were mixed with a fragment encoding an antibiotic cassette and ligated by Gibson isothermal assembly (30). The resulting fragments were transformed into PY79 selecting kanamycin and tetracycline resistance respectively. The constructs were confirmed by amplifying the region with primer pair 8486/8487 and sequencing the amplicon.

To generate the PtsG-GFP allele containing chloramphenicol (cat) resistance cassettes, the cassette was amplified from pAC225 (generous gift of Arnaud Chastanet and Rich Losick, Harvard University) with primer pair 3250/3251. Two other fragments containing the 5’ *ptsGΩpTnFLXgfp* insertion plus the region upstream and a fragment containing the 3’ end of the *ptsGΩpTnFLXgfp* insertion plus the region downstream were separately PCR amplified from DB1567 chromosomal DNA with primer pairs 8486/8884 and 8487/8885. The three fragments were mixed and ligated by Gibson isothermal assembly (30). The resulting fragment was transformed into PY79, selecting for chloramphenicol resistance, and confirmed by amplifying the region with primer pair 8486/8487 and sequencing the amplicon.

To generate chloramphenicol resistant variant fluorophores fused to PtsG, the gene for chloramphenicol (cat) resistance was amplified from pAC225 with primer pair 3250/3251. Next the upstream and downstream arms were amplified as follows: *ptsGΩpTnFLXcfp* was amplified from DB1854 chromosomal DNA with primers 8486/9043 and 8487/8885, *ptsGΩpTnFLXmvenus* was amplified from DB1855 with primers 8486/9042 and 8487/8885, and *ptsGΩpTnFLXmlychee* was amplified from DB3499 with primers 8486/9087 and 8487/8885.

The cognate arms were mixed with the chloramphenicol resistance cassette amplicon and ligated by Gibson isothermal assembly (30). The resulting fragment was transformed into PY79, selecting for chloramphenicol resistance, and confirmed by amplifying the region with primer pair 8486/8487 and sequencing for amplicon.

### iPCR and Sequencing Products

An iPCR product containing the transposed gene was amplified from *B. subtilis* chromosomal DNA of each mutated strain with the primers 6212/6420. The iPCR products were then sent for sequencing using primer 6224.

## Microfluidics

### Materials

Materials used to fabricate microfluidics devices included: glass slides (50 mm by 75 mm) from Corning, Inc.; poly(dimethylsiloxane) (PDMS, Sylgard 184) from Dow Corning, Inc.; No. 1.5 Gold Seal coverslips (48 mm by 60 mm) from VWR International, LLC; SU-8 2010 photoresist and SU-8 developer from Kayaku Advanced Materials, Inc.; titanium di-isopropoxide bis(2,4-pentanedionate) from Gelest, Inc.; and all other chemicals from Sigma-Aldrich Co.

### Device Fabrication and Operation

The microfluidic devices consisted of three layers including a glass cover slide (bottom), a ∼100-µm thick PDMS fluid layer (middle), and a 3-mm thick PDMS control layer (top), and the PDMS layers were fabricated through a mold-replica process (Baker et al. 2016; Yu et al. 2020). The SU-8 mold for the fluid layer was formed in two steps: electron beam lithography (FEI Quanta 600F scanning electron microscope equipped with a JC Nabity Nanometer Pattern Generation System) generated an array of approximately 1000 microchannels (1.0 µm deep and 0.5 to 1.0 µm wide) followed by photolithography (OAI 200 Mask Aligner) to pattern microchannels (20 µm deep and 100 µm wide) positioned perpendicularly to the microchannel array. The microchannels in the control layer were generated in a single photolithographic step.

PDMS in a 10:1 ratio of polymer to cross-linker was spin-coated onto the fluid-layer mold and poured onto the control-layer mold and degassed. The PDMS for the fluid layer was fully cured whereas the PDMS for the control layer was partially cured. The PDMS replicas were removed from the molds, and a biopsy punch was used to bore holes through the control layer. The fluid and control layers were aligned and fully cured for 4 h. The side of the fluid layer with the microchannel array was plasma cleaned (Harrick PDC-001) and bonded to the glass coverslip. The assembled device was placed in holder that was pneumatically connected to a control box to acuate on-device valves and pumps to control movement of media and cells.

Microchannels were filled with 1% bovine serum albumin (BSA) in media, and valves were checked for functionality. Media was fed through the device by gravity and directed to various locations on the device by opening and closing the valves with vacuum and pressure, respectively. Cells were loaded and trapped in the microchannel array by applying pressure above the microchannel array through the control layer. Cells were allowed to grow and acclimate for approximately 2 h before analysis began.

### Time-Lapse Image Acquisition

Fluorescence images were collected on a Nikon Ti2 microscope with a Hamamatsu ORCA-Fusion Digital CMOS camera, 100X Plan Apo oil-immersion objective (1.45 NA), and Lumencor spectra light engine. Fluorophores were excited with 474/27 nm for GFP and mNeongreen and 554/23 nm for mCherry, and their emission was collected at 515/30 nm for GFP and mNeongreen and 595/31 nm for mCherry. Semrock Triple Band Pinkel multiband and Quad Band Sedat multiband LED filter sets were coupled with single band emission filters for general fluorophore spectra. The objective warmer (Okolab) was set at 30°C for all experiments. Nikon NIS-Elements was used for multi-color image acquisition at 2-min intervals at 12 locations within the microchannel array. Denoise.ai mode was used to align images due to stage jitter.

### Data Analysis

Image files in ND2 format were parsed into 12 separate files based on the 12 locations imaged within the microchannel array. With Fiji software (ImageJ), horizontal positions of the images were adjusted to 0° to track cell position over time, and backgrounds were subtracted from each set of images. Kymographs of individual channels containing viable cells were generated from images taken at 2-min intervals, and localization of PtsG-GFP (membrane), mCherry or mNeongreen (FtsZ), and mCherry (cytoplasm) were manually tracked across the kymographs. To track cell division, cell length, and FtsZ ring dynamics, a rectangular region-of-interest for each cell was used to integrate the fluorescence signals and generate single-line profiles parallel to the cell length. For the cytoplasmic dye (mCherry), a 20% local decrease between adjacent cells was used as the threshold to indicate cytokinesis, whereas for the membrane label (PtsG-GFP), a maximum in the fluorescence signal was used as the threshold for cytokinesis. Progeny of the mother cells were followed by continuous tracking of fluorescence signals.

### Fluorescent Activated Cell Sorting

Sorting was done by the Indiana University Bloomington (IUB) Flow Cytometry Core Facility (FCCF, RRID: SCR_024398). Samples were grown in LB broth overnight at room temperature rolling. The next day, 100 µl of culture was back diluted into 3 ml of LB broth and rolled at 37°C for 1 h or until the OD_600_ reached 0.3. 1 ml of cells were collected in a 2 ml Eppendorf tube and centrifuged at 14,000 RPM for 2 min. The supernatant was removed, and the pellet resuspended in 1 ml of 1x PBS buffer. Samples were then taken to the IUB FCCF for sorting based on the scatter and fluorescence properties of the cells.

The instrument used for the sort was a 5-laser SORP FACSAriaIIu with FACSDiva software version 9.4 (BD Biosciences, San Jose, CA). WT DK1042 cells, which have no inherent fluorescence, were used to set the boundary between msfGFP negative and positive cells. Cells were sorted at room temperature with an 85-micron nozzle, set at 35 psi, in 4-way purity sort mode. A 488 nm 100 mW laser was used for scatter and msfGFP detection. The parameters used for the sorting were FSC-PMT at 300 V, SSC at 269 V, and B530/30 at 650 V, all on log-scaled axes. To exclude instrument noise, debris and other non-bacterial particles, the threshold was set at 200 on SSC. 3,800 fluorescent cells were collected in LB broth and grown up rolling overnight at 37°C. Cells were spun down in glycerol and stored at – 80°C for further analysis.

## ACKNOWLEDGEMENTS

We thank Laura Lastra for the construction of pLL4. We thank Christiane Hassel of the Indiana University Bloomington Flow Cytometry Core facility for technical support and the IU Nanoscience Core Facility for use of its instruments. This work was funded in part by NIH GM131783 to DBK and NIH R35GM141922 to SCJ. Additional support was provided from the NIH Graduate Training Program in Quantitative and Chemical Biology under Award Number NIH T32GM131994 and Indiana University to JHJ.

